## Supplementary figures and images for "Phylogenomics investigation of sparids (Teleostei: Spariformes) using high-quality proteomes highlights the importance of taxon sampling"

### Supplementary Figure 1

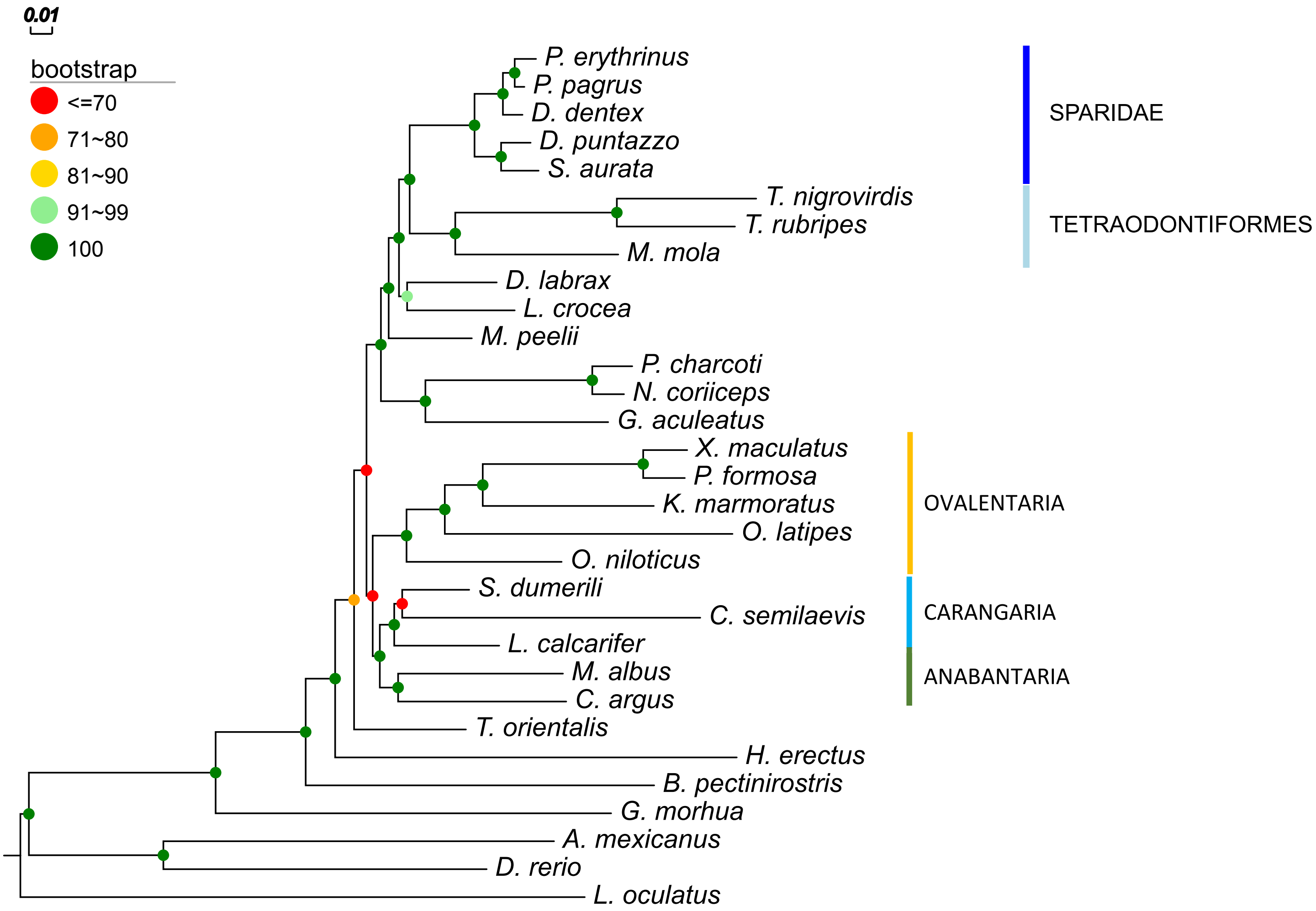

### Supplementary Figure 2A

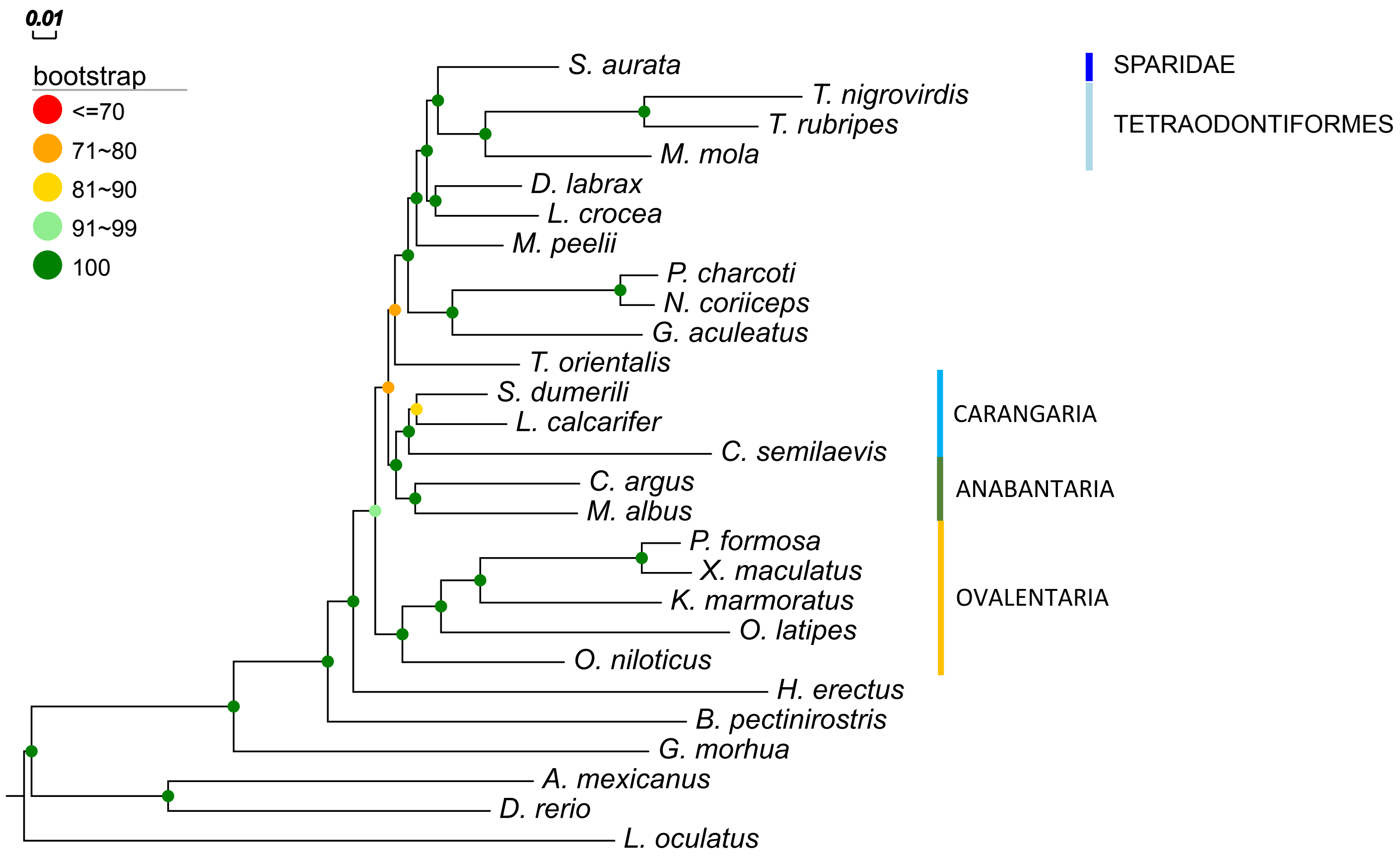

### Supplementary Figure 2B

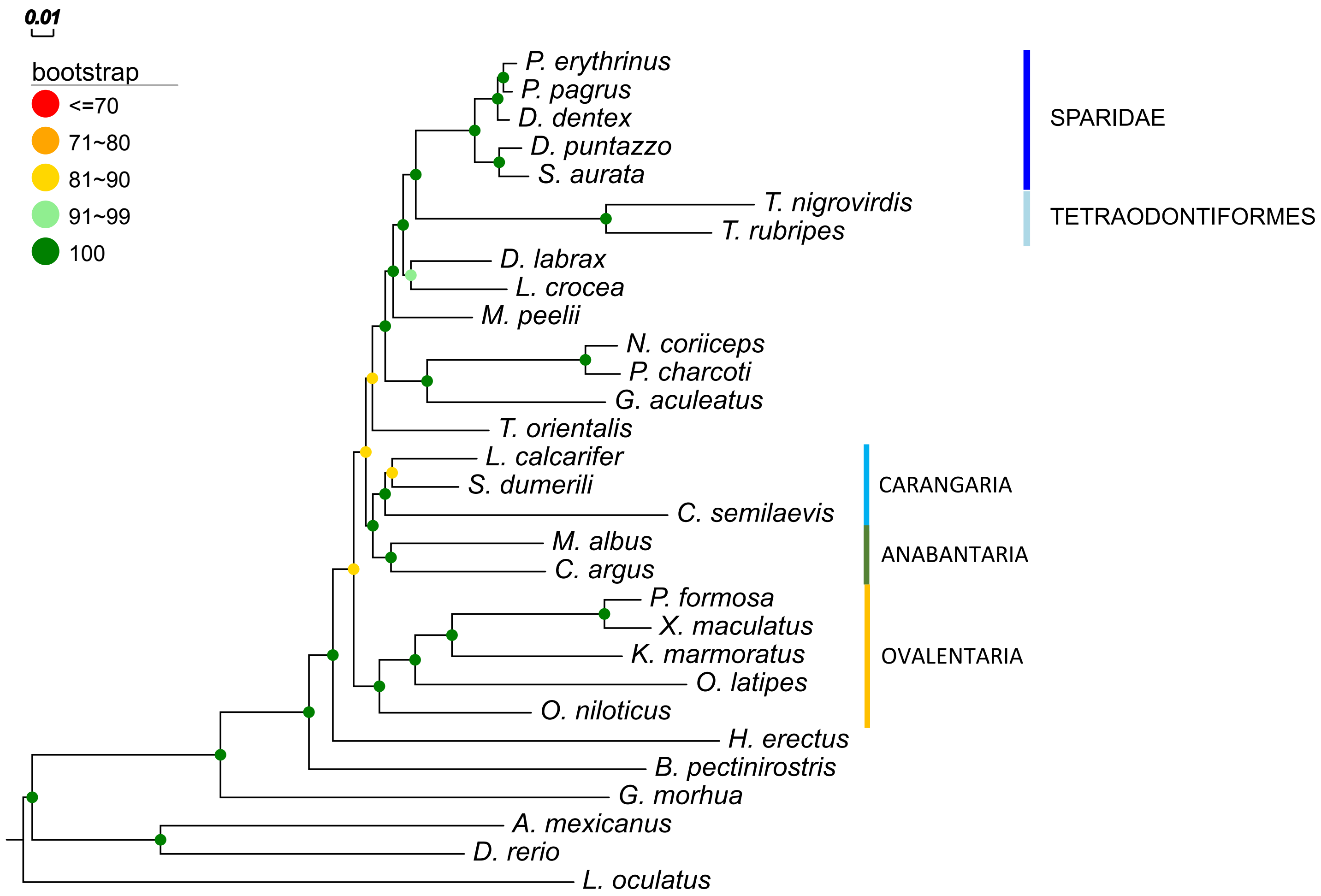

### Supplementary Figure 2C

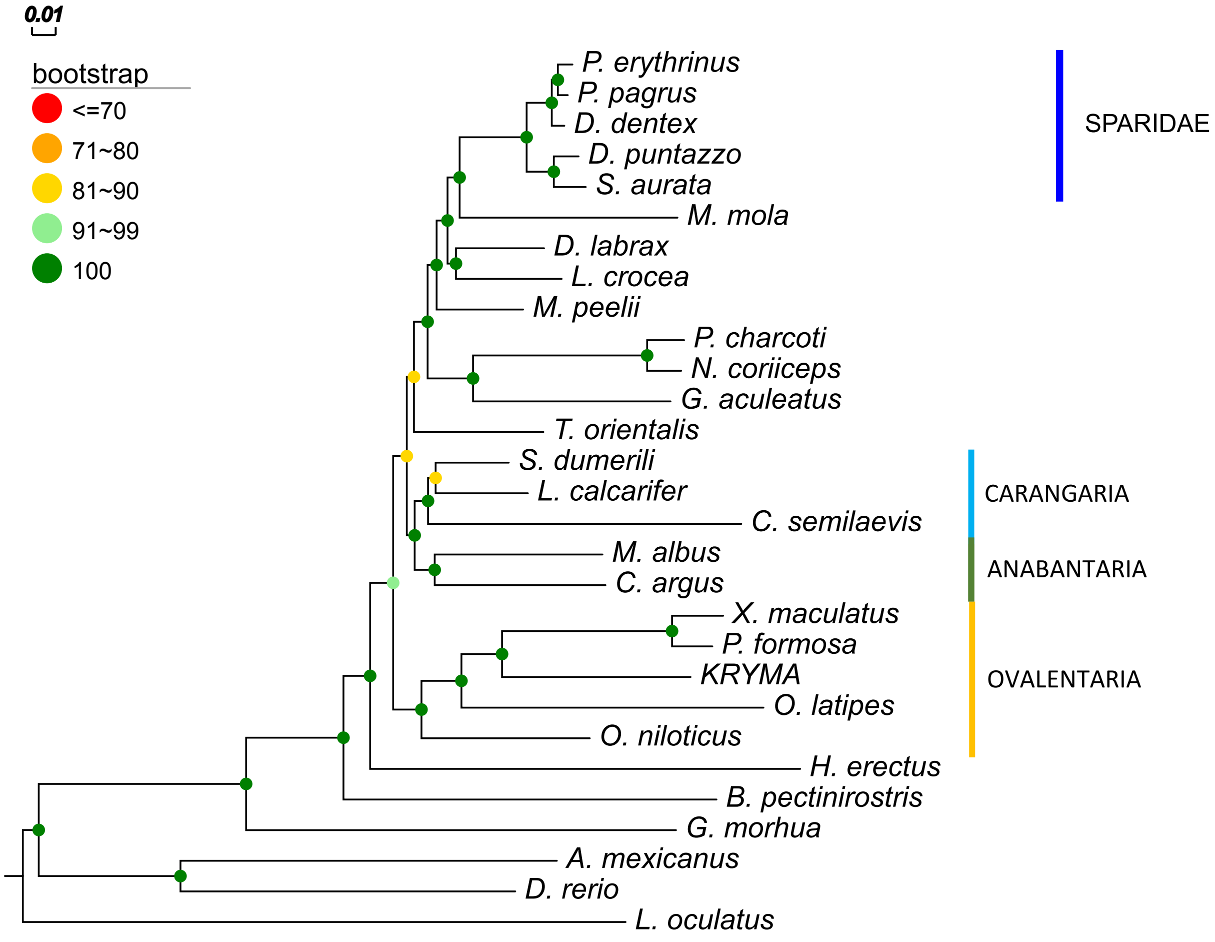

### Supplementary Figure 3A

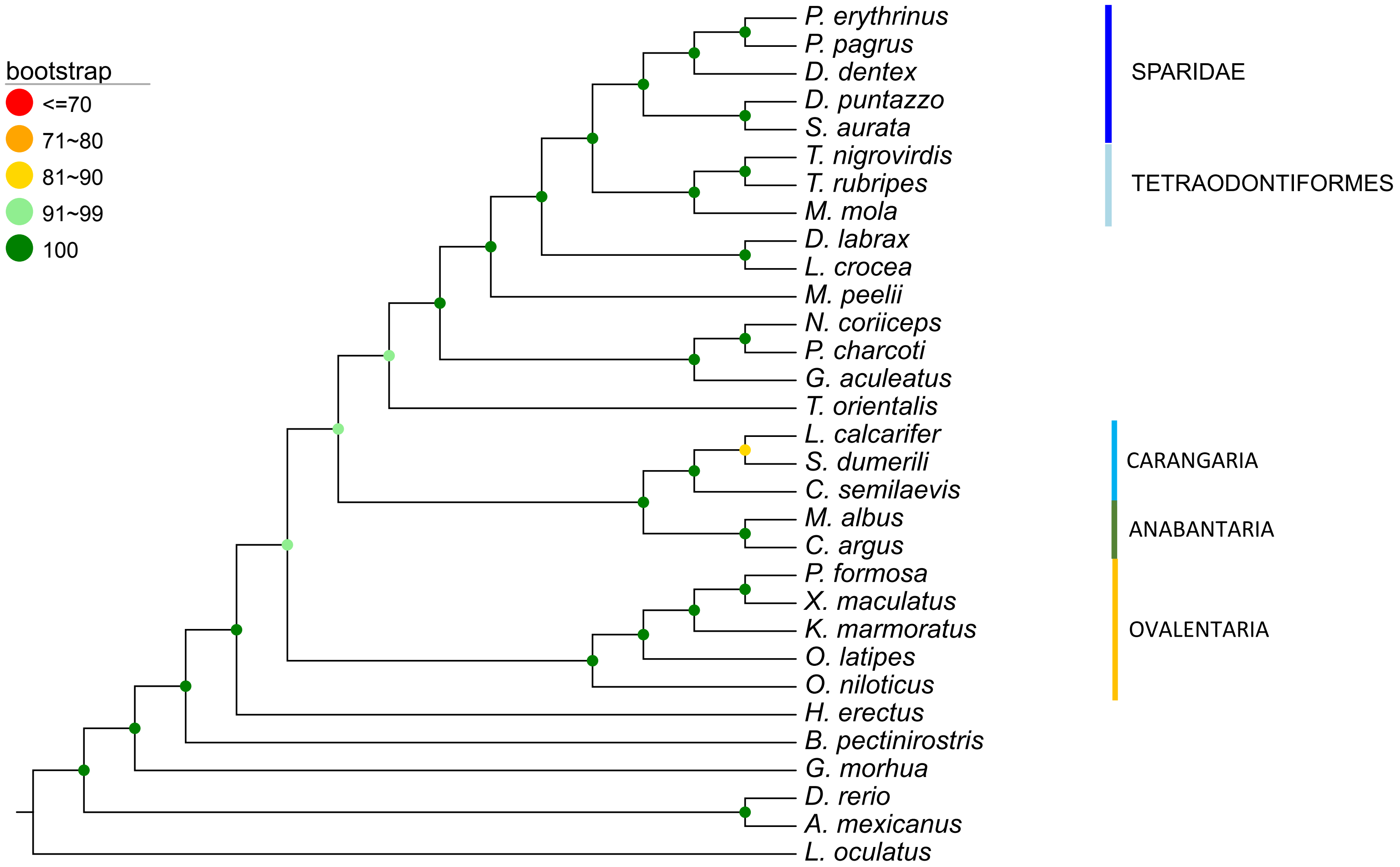

### Supplementary Figure 3B

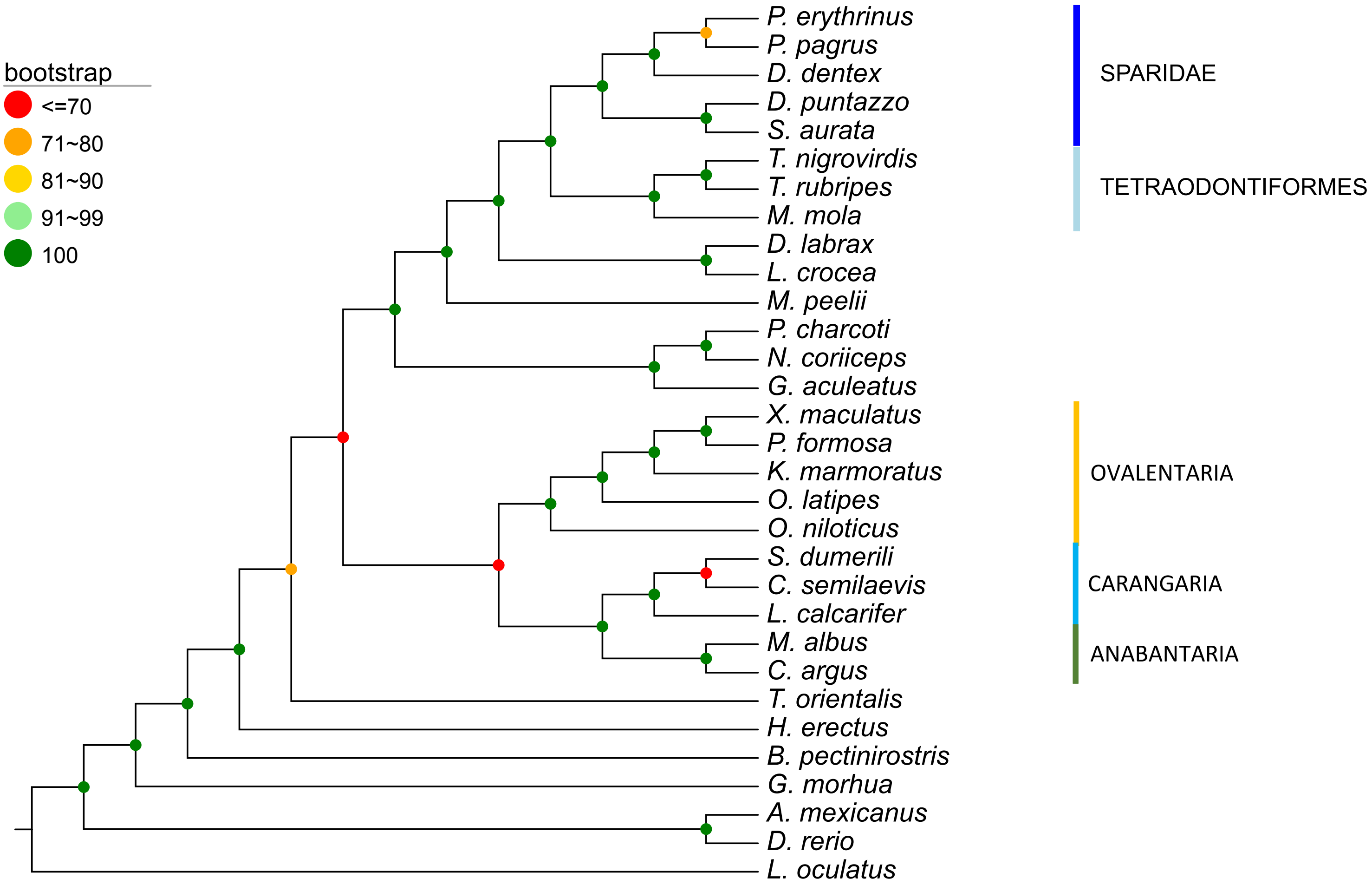

### Supplementary Figure 4A

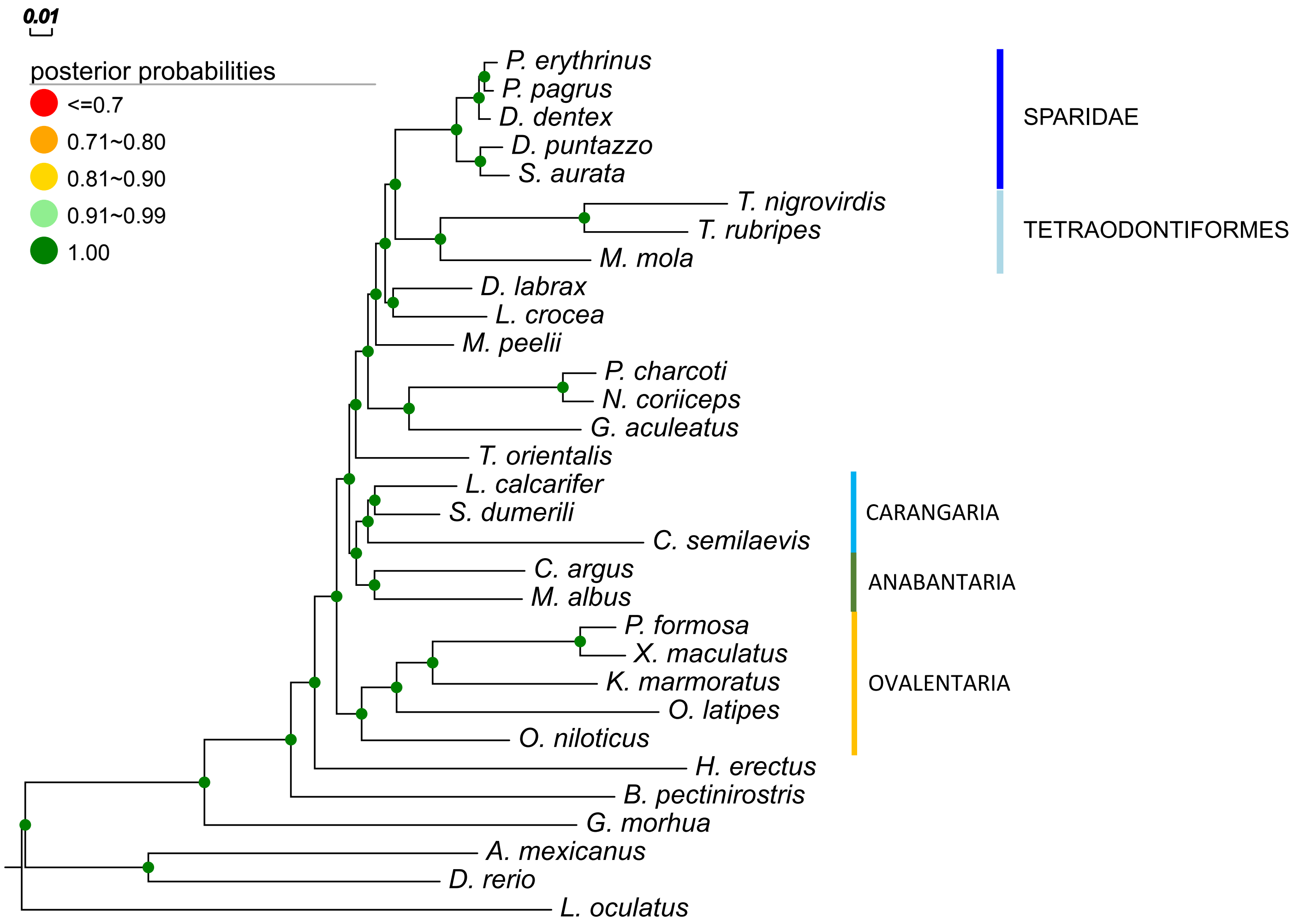

### Supplementary Figure 4B

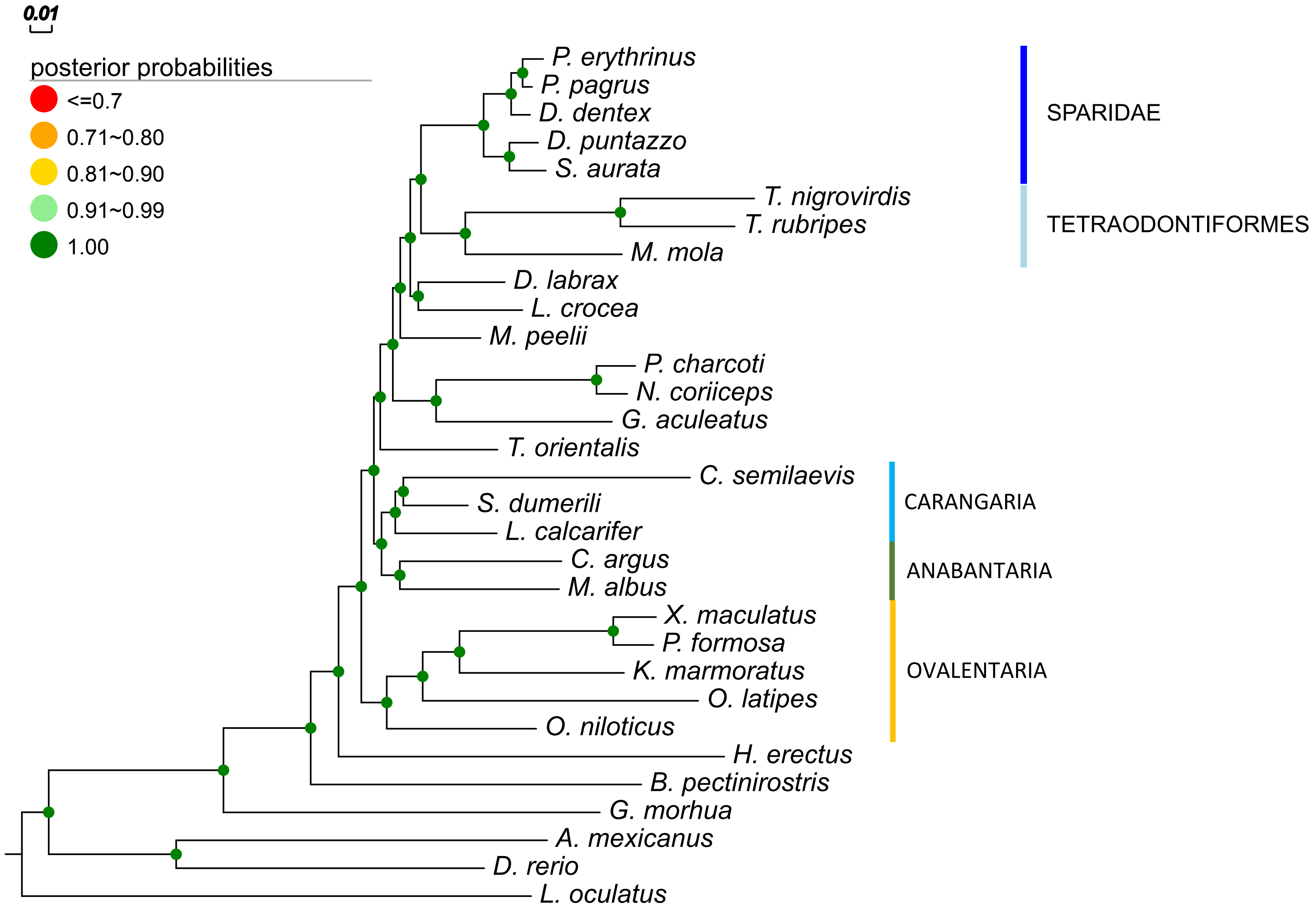

### Supplementary Figure 5A

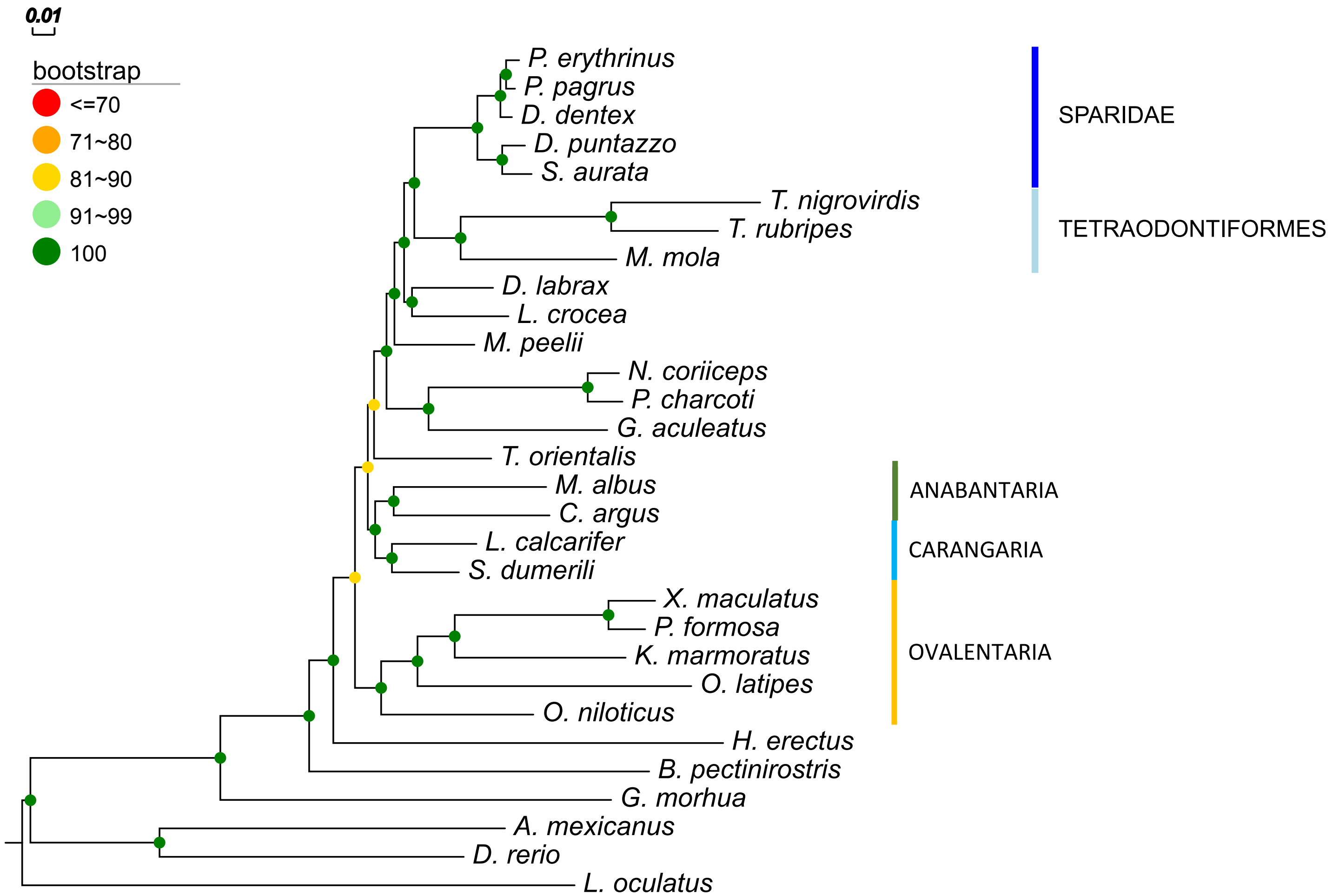

### Supplementary Figure 5B

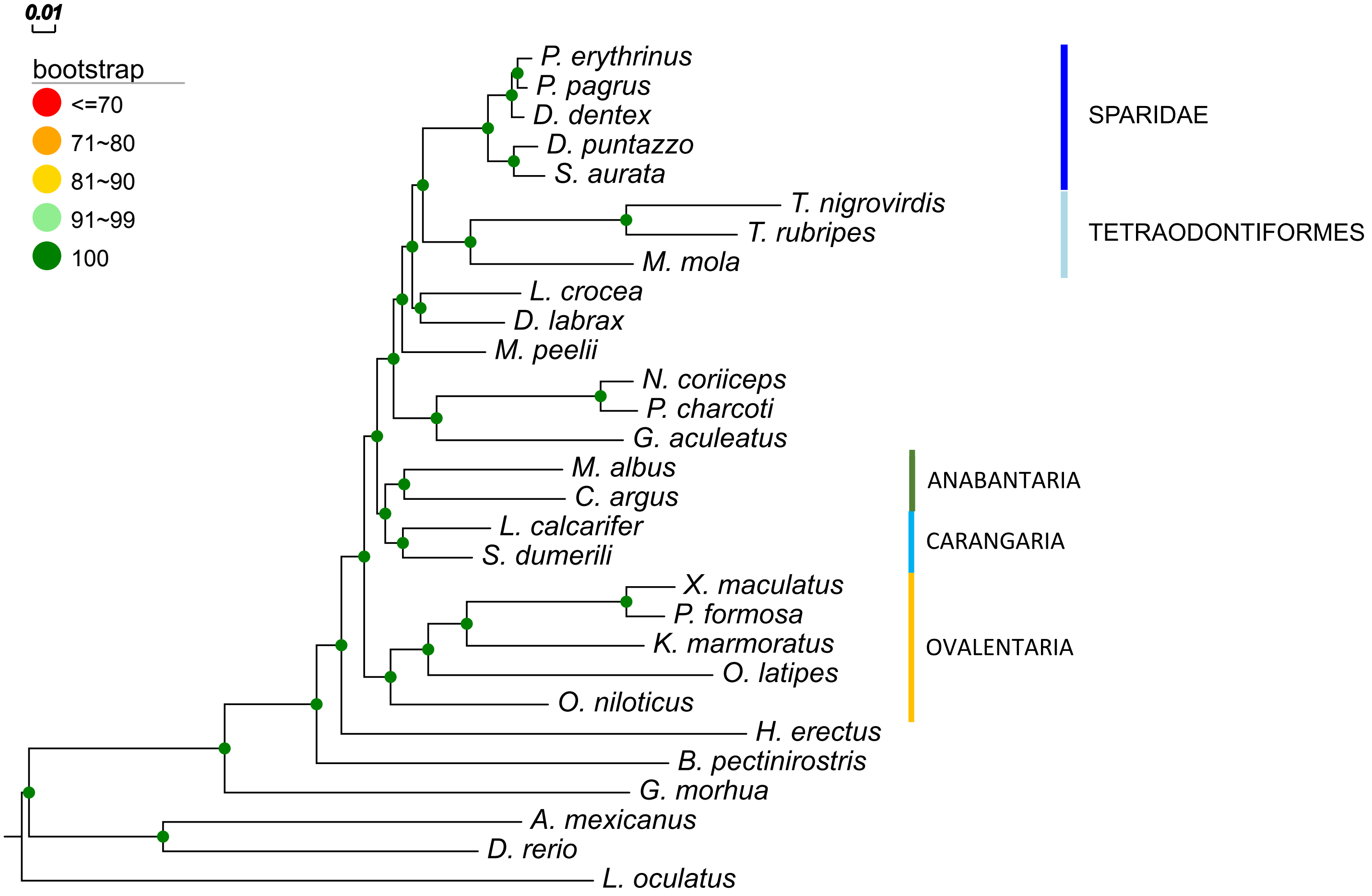

### Supplementary Figure 6A5

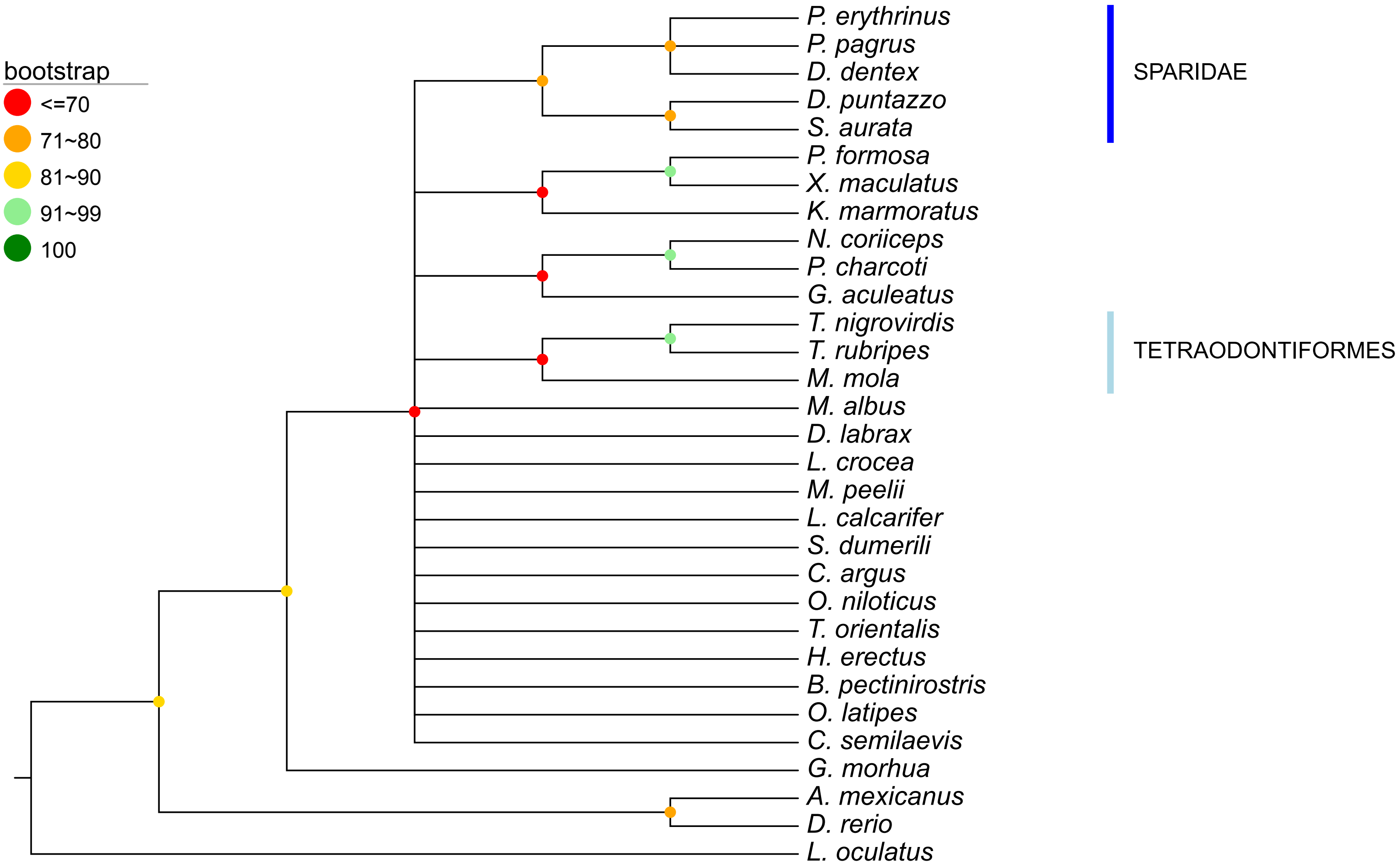

### Supplementary Figure 6B

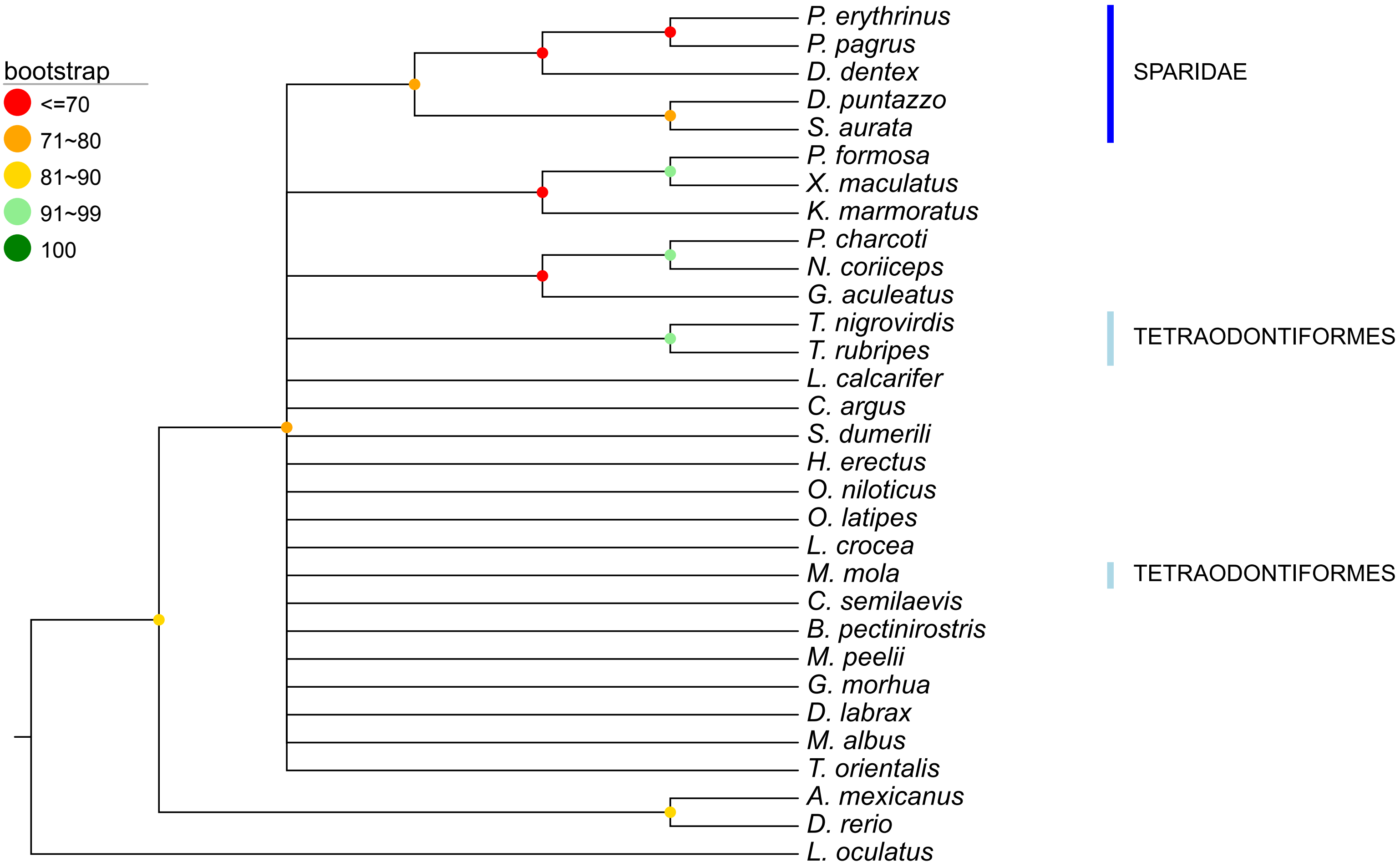

### Supplementary Figure 6C

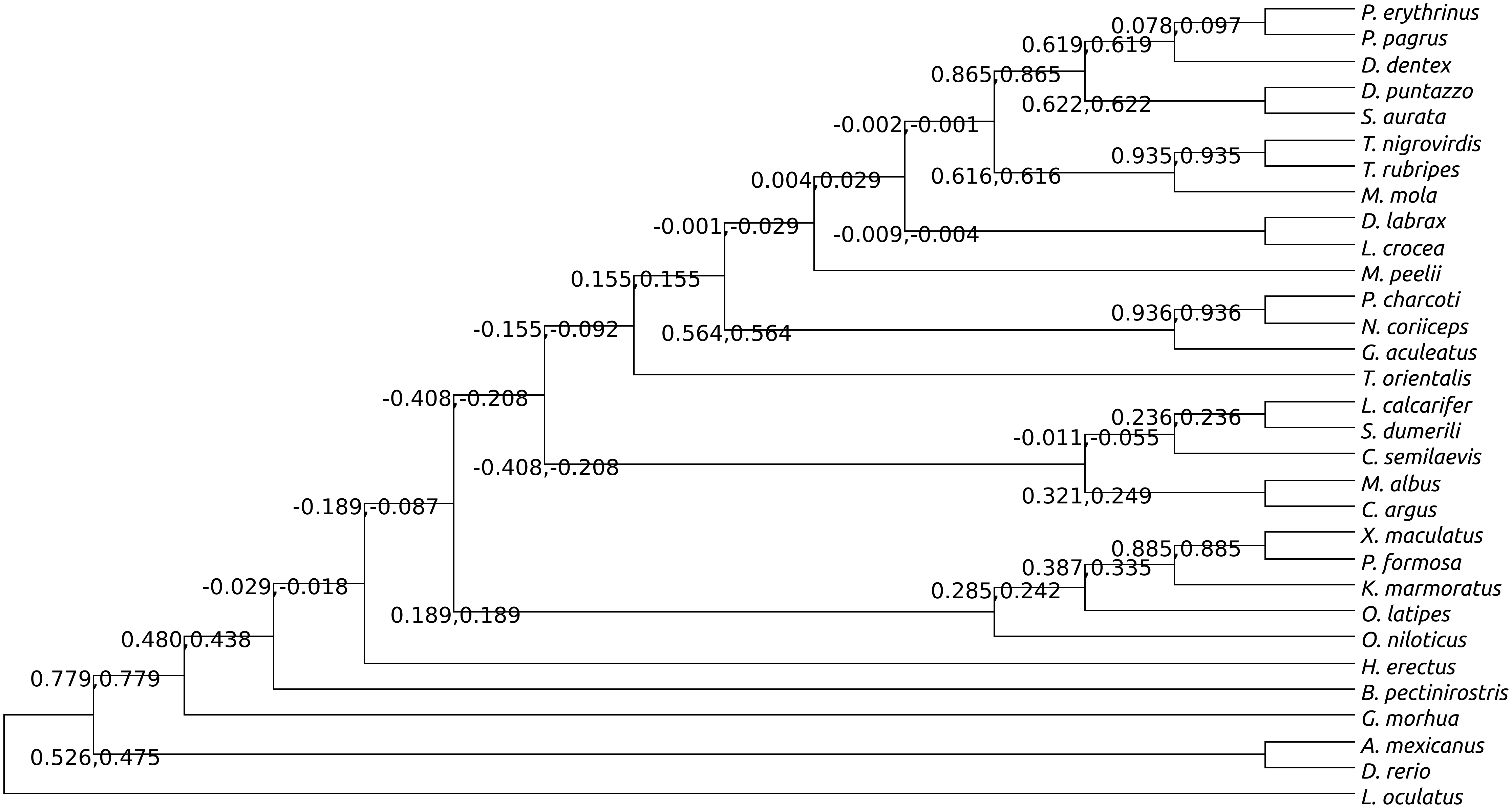

### Supplementary Figure 6D

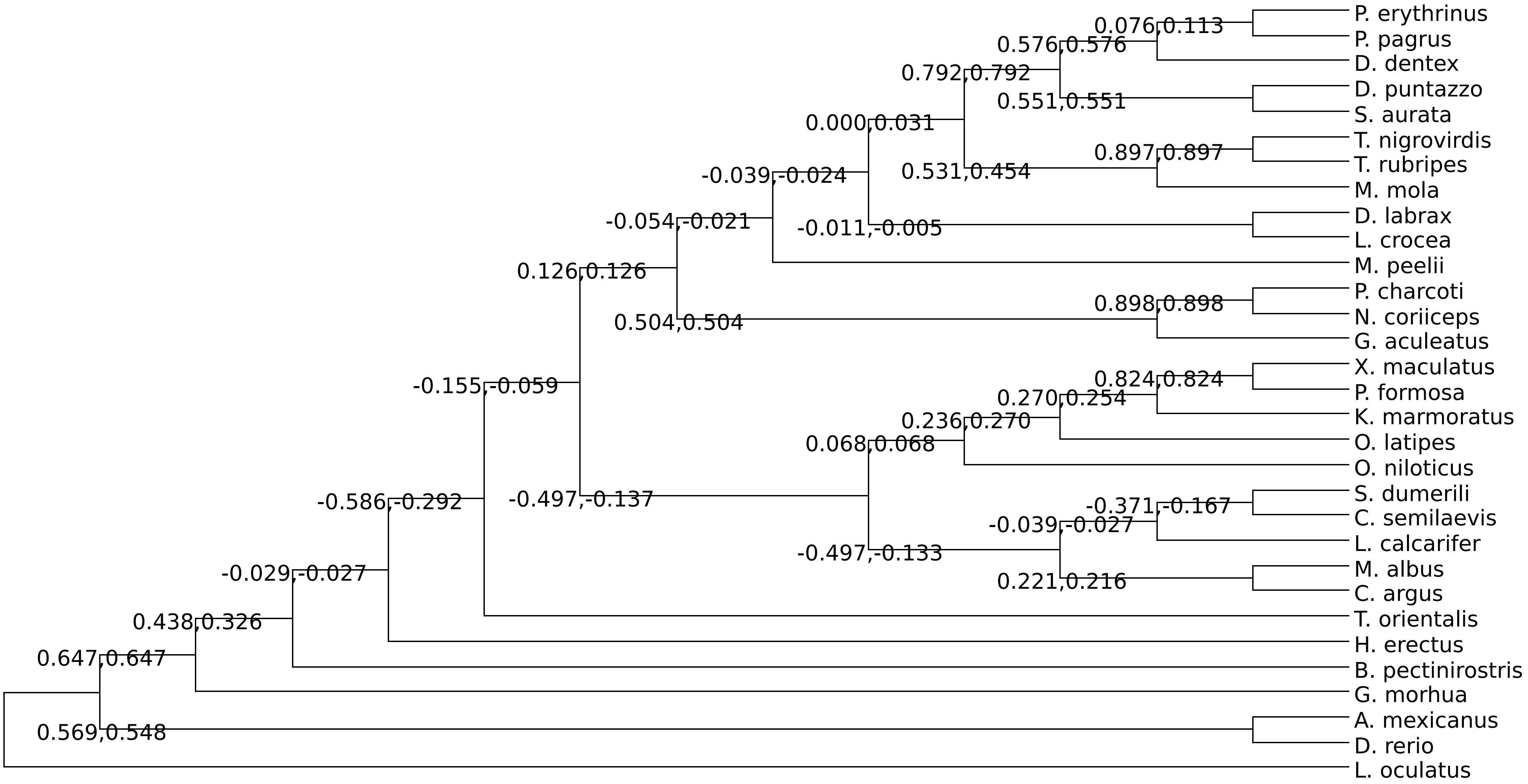

### Supplementary Figure 7A

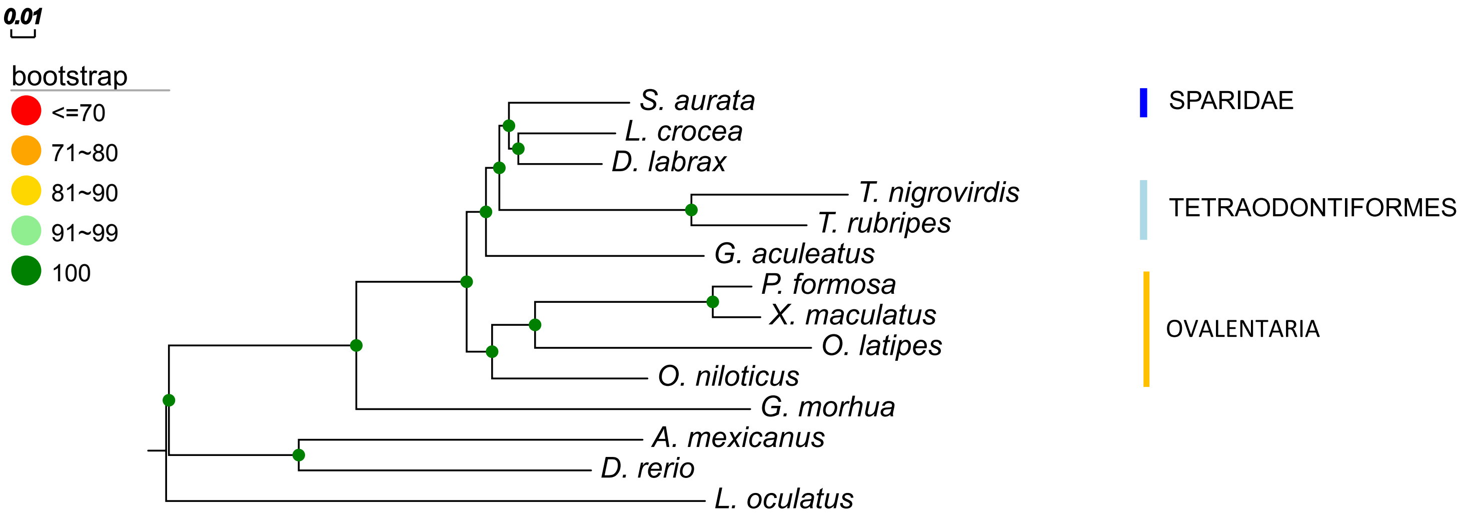

### Supplementary Figure 7B

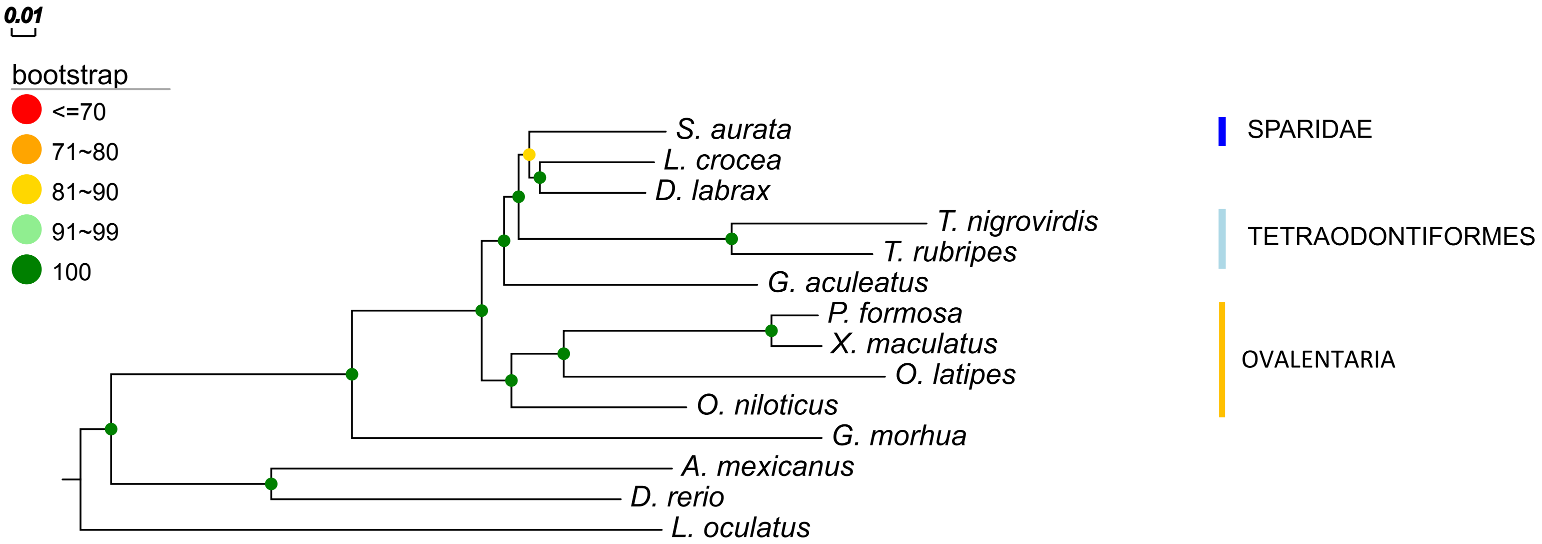
